# BoLA-DRB3 gene haplotypes show divergence in native Sudanese cattle from Taurine and Zebu breeds

**DOI:** 10.1101/2020.08.07.241133

**Authors:** Bashir Salim, Shin-nosuke Takeshima, Ryo Nakao, Mohamed A-M Moustafa, Mohamed-Khair A. Ahmed, Sumaya Y. Kambal, Joram M. Mwacharo, Guillermo Giovambattista

## Abstract

Autochthonous Sudanese cattle breeds, namely Baggara for beef and Butana and Kenana for dairy, are characterized by their adaptive characteristics and high performance in hot and dry agro-ecosystems, are used largely by nomadic and semi-nomadic pastoralists. Here we analyzed the diversity and genetic structure of the BoLA-DRB3 gene, a genetic locus linked to the immune response, for the indigenous cattle of Sudan and in the context of the global cattle repository. Blood samples (n=225) were taken from three indigenous breeds (Baggara; n=113, Butana; n= 60 and Kenana; n=52) distributed across six regions of Sudan. Nucleotide sequences were genotyped using the sequence-based typing method. Sequence electropherograms were analyzed using the Assign SBT software. We describe 53 alleles, including seven new, novel alleles. In the Baggara breed the number of alleles was 46 (40 previously reported and six new ones), 33 in the Kenana breed (28 previously reported and five new ones), and 33 in the Butana breed (28 previously reported and five new ones). Venn analysis of Sudanese breeds with Southeast Asian, European and American cattle showed 115 alleles of which 14 were unique to Sudanese breeds. Three of the alleles exhibited gene frequency of > 0.5%, representing 26% of the 53 alleles detected in the native Sudanese cattle. Observed versus expected heterozygosity was higher than 0.93 in all three breeds analyzed and equilibrium status revealed by Hardy-Weinberg Equilibrium suggests pure genetic drift. Gene frequency distributions of Baggara cattle showed an even distribution (*P* = 0.016), consistent with the theoretical proportion expected under balancing selection pressure as opposed to positive or neutral selection. In contrast, Butana and Kenana cattle (*P* = 0.225 and *P* = 0.138, respectively) were more congruent with neutral selection, similar to the results obtained for most of the cattle breeds analyzed so far. Sudanese cattle breeds were located within a narrow cloud in an intermediate position between the Zebu and Taurine breeds and close to other Southeast Asian breeds, in accordance with the composite origin of these native breeds, which is also reinforced by the presence of African and Zebu unique BoLA-DRB3 alleles within these breeds. The results of the Principal Component Analysis were in agreement with the overall clustering pattern observed on the NJ and/or UPGMA trees. These results contribute to our understanding of the genetic diversity and distribution pattern of BoLA-DRB3 gene alleles in Sudanese cattle breeds and provide insight into their uniqueness in their ability to survive arrays of tropical diseases and reproduce well in Sudan’s harsh environment.

**Author summary:** African cattle survive and adapt to a variety of diseases via acquired immunity capable of presenting antigens through the function of the major histocompatibility complex (MHC) or bovine leukocyte antigen (BoLA) in cattle. The aim of this study was to investigate how the immune system is structured and to what extent three economically important breeds in Sudan differ from exotic cattle. Here, we use the sequence-based typing approach to analyze BoLA-DRB3’s genetic diversity linked to immunity against complex diseases that infect cattle. By examining 225 indigenous cattle belonging to three breeds in Sudan, we demonstrate that these cattle are unique from all known cattle by identifying seven new alleles; *BoLA-DRB3*\**004:02Sp, BoLA-DRB3*\**011:02Sp, BoLA-DRB3*\**018:01Sp, BoLA-DRB3*\**021:01sp, BoLA-DRB3*\**024:18Sp, BoLA-DRB3*\**027:05sp*, and *BoLA-DRB3*\**032:01sp*. When analyzing frequency of the protein pockets implicated in the antigen-binding function of the MHC complex by PCA we found that pockets 4 and 9 are the ones that best differentiate these native breeds from the rest. This may be attributed to high disease tolerance/susceptibility to tropical infections, such as those carried by ticks and intestinal parasites. Further studies are needed on these newly identified variants and their association with specific common disease(s). This finding is especially important for disease resistance/susceptibility association to help advise on candidate animals in selection schemes.

## Introduction

There is a consensus among population geneticists that the Sudanese cattle populations belong to the humped Zebu cattle breed and are classified into two principal varieties; northern Sudan and Nilotic [1,2]. The Kenana and Butana breeds are the best known milk-producing northern Sudan Zebu breeds [3–5] with yield amounts of more than 1,500kg of milk per lactation [6–8]. Phenotypically, the Kenana breed, that is predominantly found in the Blue Nile state, is distinguished by a light blue-gray coat color, darker at the hooves and head, in contrast to the red-coated cattle of the Batahin and Shukria tribes inhabiting the desert area between the Blue Nile and the River Atbara, named the Butana breed [1]. On the other hand, the Baggara breed is the major fattening northern Sudan Zebu cattle breed that is raised by Baggara Bedouin pastoralists, hence the name, and is concentrated chiefly in the western region of Sudan, Niger, Chad, Cameroon and Nigeria. They are characterized by short horns and a large hump and are found particularly in the Darfur and Kordofan regions where they display white or white with black markings (Nyalawi population), and red or dark red (Daeinawi Aka Messairi/Rezaigi population; [9]).

Genetic factors affecting disease resistance or susceptibility determine cattle wellbeing and reproduction. The immune system in vertebrates is an adaptive defense mechanism which has evolved to defend them from invasive pathogens [10]. The genes of the Major Histocompatibility Complex (MHC) are becoming more popular among animal breeders, veterinary geneticists and immunologists as they are associated with genetic resistance and susceptibility to a wide variety of diseases [11]. Genetic characterization of MHC polymorphism can inform the design of breeding programs to reduce the manifestation and severity of infectious diseases, especially in domestic animals and cattle breeding schemes [12]. The association of MHC with diseases in ruminants is well documented [13–18]. MHC gene genome organization is assigned to Bovine Chromosome 23 (BTA 23) [13,14] and it is called Bovine Leukocyte Antigen (BoLA). Recently, Kim *et al*., [15] investigated five African breeds for the identification of common and unique African genome-specific selection signatures and compared them with commercial breeds. In this comparison, when analyzing in depth the bovine lymphocyte antigen (BoLA) region, six BoLA haplotype blocks were identified and the major African cattle haplotypes correspond to minor haplotypes in commercial cattle. Detailed genomic analysis of the BoLA is important due to the crucial role it plays in an animal’s immune system. The BoLA molecules’ extensive structural polymorphism is responsible for differences in the immune response to infectious agents among animals. For instance, high polymorphism observed at the DRB3 locus can help to recognize superior haplotypes for tick infestation resistance [15]. MHC research may also assist in the formation and design of synthetic peptide-based vaccines containing one or more pathogen T-cell epitope.

Despite its central role in the immune response, the genetic diversity of the BoLA-DRB3 gene has been characterized in a limited number of breeds and cross-breeds from Europe, Asia and the Americas using polymerase chain reaction-sequence based typing (PCR-SBT; [16–29]) and target next generation sequencing (Target-NGS; [30]), the most powerful tools used to identify diversity of BoLA-DRB3 alleles in cattle breeds. Until now, private African BoLA-DRB3 alleles have been reported by authors using indirect techniques, such as PCR-RFLP, followed by cloning and sequencing [31–34]. These studies focused mainly on screening and analysis of a few animals from some African breeds (e.g., Sanga, Kenana, Butana); however, to the best of our knowledge, no study has used SBT or NGS assays to examine genetic diversity of BoLA-DRB3 in large samples of native African cattle breeds. Previous work showed the presence of a high number of private alleles in native breeds. Consequently, there are still a number of breeds that remain uncharacterized, and this number only increases when local native bovine breeds are considered [18,26,27,29,30].

This study was undertaken to examine patterns of genetic variation of BoLA-DRB3 alleles in Baggara, Butana and Kenana native cattle breeds of Sudan and to compare these with commercial breeds, specifically, 1) to identify any unique/novel alleles for Sudanese native cattle breeds and alleles that confer tick resistance, and 2) to make available genetic information on resistance to infectious diseases that can be used as a basis for genetic evaluation and for designing appropriate breeding schemes.

## Results

### Distribution of *BoLA-DRB3* alleles in selected native Sudanese cattle breeds

Polymerase chain reaction-sequence-based typing (PCR-SBT) genotyping allowed us to identify 53 *BoLA-DRB3* alleles (46 previously reported variants and seven new alleles; Table 1) for the native breeds selected in this study. The number of alleles (n_a)_ was 46 in Baggara cattle (40 previously reported and six new), 33 in Kenana cattle (28 previously reported and five new), and 33 in Butana cattle (28 previously reported and five news) (Tables 1 and 2a). The new BoLA-DRB3 variants were confirmed by the presence of at least three carrier animals and in two breeds for each of them. Nucleotide and predicted amino acid sequences of the seven new allele variants are shown in Fig. 1.

**Table 1.**
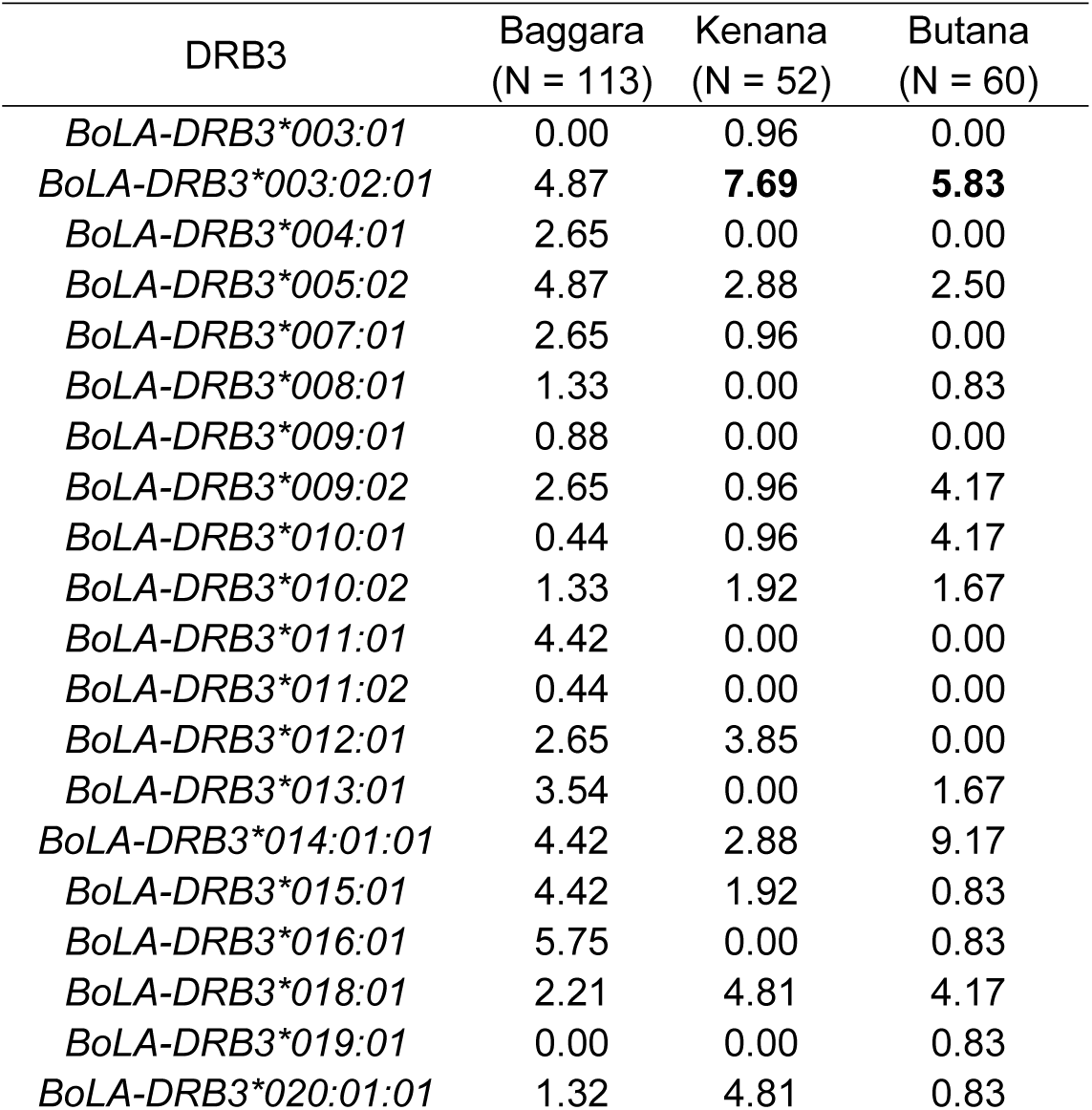
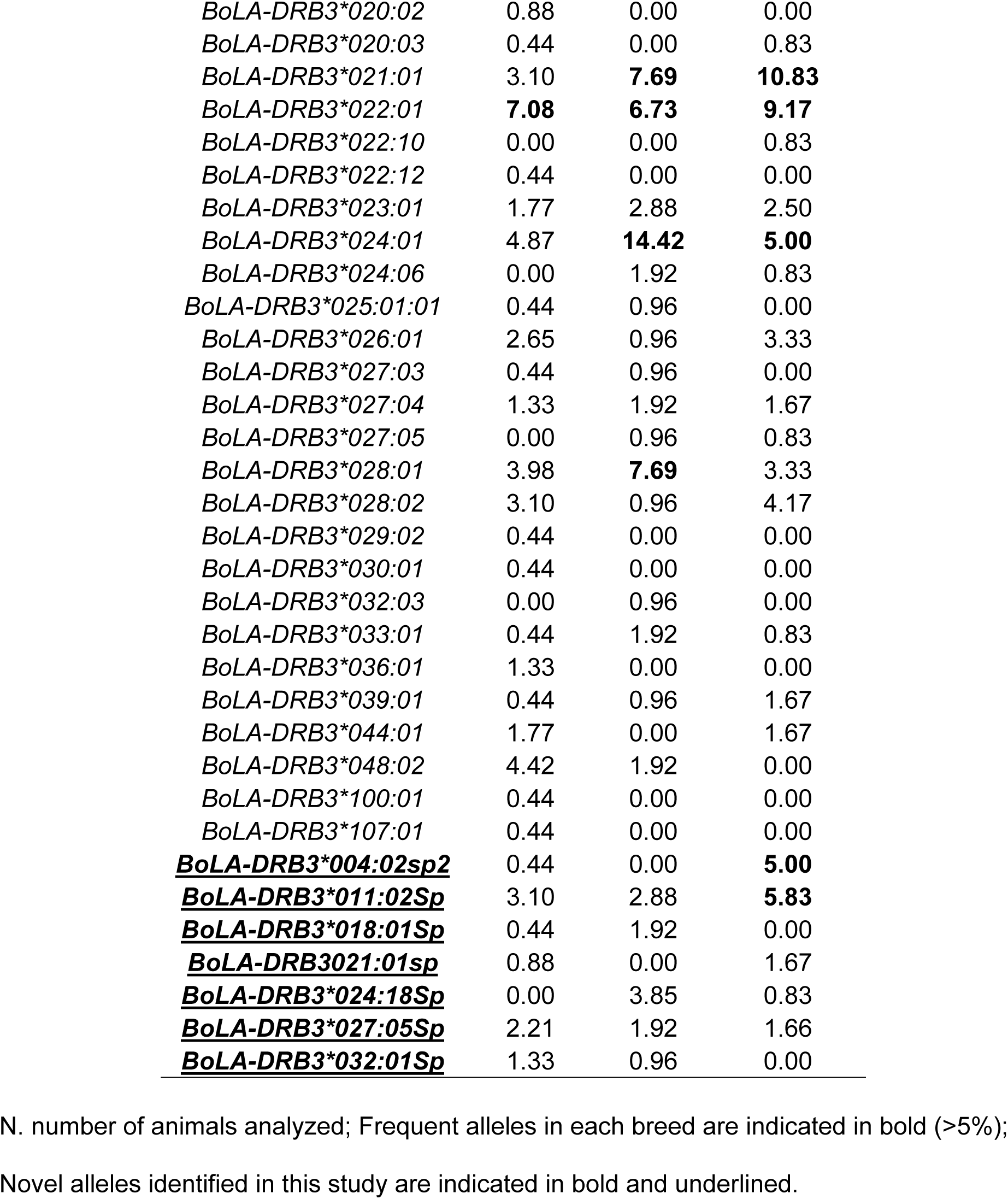
BoLA-DRB3 allele frequencies in native Sudanese cattle breeds

| DRB3 | Baggara<br>(N = 113) | Kenana<br>(N = 52) | Butana<br>(N = 60) |
| --- | --- | --- | --- |
| *BoLA-DRB3\*003:01* | 0.00 | 0.96 | 0.00 |
| *BoLA-DRB3\*003:02:01* | 4.87 | **7.69** | **5.83** |
| *BoLA-DRB3\*004:01* | 2.65 | 0.00 | 0.00 |
| *BoLA-DRB3\*005:02* | 4.87 | 2.88 | 2.50 |
| *BoLA-DRB3\*007:01* | 2.65 | 0.96 | 0.00 |
| *BoLA-DRB3\*008:01* | 1.33 | 0.00 | 0.83 |
| *BoLA-DRB3\*009:01* | 0.88 | 0.00 | 0.00 |
| *BoLA-DRB3\*009:02* | 2.65 | 0.96 | 4.17 |
| *BoLA-DRB3\*010:01* | 0.44 | 0.96 | 4.17 |
| *BoLA-DRB3\*010:02* | 1.33 | 1.92 | 1.67 |
| *BoLA-DRB3\*011:01* | 4.42 | 0.00 | 0.00 |
| *BoLA-DRB3\*011:02* | 0.44 | 0.00 | 0.00 |
| *BoLA-DRB3\*012:01* | 2.65 | 3.85 | 0.00 |
| *BoLA-DRB3\*013:01* | 3.54 | 0.00 | 1.67 |
| *BoLA-DRB3\*014:01:01* | 4.42 | 2.88 | 9.17 |
| *BoLA-DRB3\*015:01* | 4.42 | 1.92 | 0.83 |
| *BoLA-DRB3\*016:01* | 5.75 | 0.00 | 0.83 |
| *BoLA-DRB3\*018:01* | 2.21 | 4.81 | 4.17 |
| *BoLA-DRB3\*019:01* | 0.00 | 0.00 | 0.83 |
| *BoLA-DRB3\*020:01:01* | 1.32 | 4.81 | 0.83 |
| *BoLA-DRB3\*020:02* | 0.88 | 0.00 | 0.00 |
| *BoLA-DRB3\*020:03* | 0.44 | 0.00 | 0.83 |
| *BoLA-DRB3\*021:01* | 3.10 | **7.69** | **10.83** |
| *BoLA-DRB3\*022:01* | **7.08** | **6.73** | **9.17** |
| *BoLA-DRB3\*022:10* | 0.00 | 0.00 | 0.83 |
| *BoLA-DRB3\*022:12* | 0.44 | 0.00 | 0.00 |
| *BoLA-DRB3\*023:01* | 1.77 | 2.88 | 2.50 |
| *BoLA-DRB3\*024:01* | 4.87 | **14.42** | **5.00** |
| *BoLA-DRB3\*024:06* | 0.00 | 1.92 | 0.83 |
| *BoLA-DRB3\*025:01:01* | 0.44 | 0.96 | 0.00 |
| *BoLA-DRB3\*026:01* | 2.65 | 0.96 | 3.33 |
| *BoLA-DRB3\*027:03* | 0.44 | 0.96 | 0.00 |
| *BoLA-DRB3\*027:04* | 1.33 | 1.92 | 1.67 |
| *BoLA-DRB3\*027:05* | 0.00 | 0.96 | 0.83 |
| *BoLA-DRB3\*028:01* | 3.98 | **7.69** | 3.33 |
| *BoLA-DRB3\*028:02* | 3.10 | 0.96 | 4.17 |
| *BoLA-DRB3\*029:02* | 0.44 | 0.00 | 0.00 |
| *BoLA-DRB3\*030:01* | 0.44 | 0.00 | 0.00 |
| *BoLA-DRB3\*032:03* | 0.00 | 0.96 | 0.00 |
| *BoLA-DRB3\*033:01* | 0.44 | 1.92 | 0.83 |
| *BoLA-DRB3\*036:01* | 1.33 | 0.00 | 0.00 |
| *BoLA-DRB3\*039:01* | 0.44 | 0.96 | 1.67 |
| *BoLA-DRB3\*044:01* | 1.77 | 0.00 | 1.67 |
| *BoLA-DRB3\*048:02* | 4.42 | 1.92 | 0.00 |
| *BoLA-DRB3\*100:01* | 0.44 | 0.00 | 0.00 |
| *BoLA-DRB3\*107:01* | 0.44 | 0.00 | 0.00 |
| **<u>BoLA-DRB3\*004:02sp2</u>** | 0.44 | 0.00 | **5.00** |
| **<u>BoLA-DRB3\*011:02Sp</u>** | 3.10 | 2.88 | **5.83** |
| **<u>BoLA-DRB3\*018:01Sp</u>** | 0.44 | 1.92 | 0.00 |
| **<u>BoLA-DRB3021:01sp</u>** | 0.88 | 0.00 | 1.67 |
| **<u>BoLA-DRB3\*024:18Sp</u>** | 0.00 | 3.85 | 0.83 |
| **<u>BoLA-DRB3\*027:05Sp</u>** | 2.21 | 1.92 | 1.66 |
| **<u>BoLA-DRB3\*032:01Sp</u>** | 1.33 | 0.96 | 0.00 |

N. number of animals analyzed; Frequent alleles in each breed are indicated in bold (>5%);
Novel alleles identified in this study are indicated in bold and underlined.

**Table 2.** Values of genetic diversity estimated for the BoLA-DRB3 in the Sudan native breeds analyzed.

| Breed | N | $n_a$ | $h_o$ | $h_e$ | HWE | Slatkin's |
| --- | --- | --- | --- | --- | --- | --- |
| | | | | | $F_{IS}$ - p value | p value |
| Baggara | 113 | 46 | 0.938 | 0.969 | 0.032 - 0.277 | **0.016** |
| Butana | 52 | 33 | 0.961 | 0.951 | 0.005 - 0.100 | 0.255 |
| Kenana | 60 | 33 | 0.950 | 0.954 | -0.011 - 0.628 | 0.138 |

| Breed | $\pi$ | NPD | Total | | | ABS | | |
| --- | --- | --- | --- | --- | --- | --- | --- | --- |
| | | | $d_s$ | $d_n$ | $d_n / d_s$ | $d_s$ | $d_n$ | $d_n / d_s$ |
| Baggara | 0.080 | 19.36 | 0.037 | 0.097 | 2.62 | 0.118 | 0.394 | 3.34 |
| Butana | 0.074 | 17.99 | 0.035 | 0.097 | 2.77 | 0.120 | 0.404 | 3.37 |
| Kenana | 0.075 | 18.28 | 0.034 | 0.096 | 2.88 | 0.115 | 0.402 | 3.50 |

**Fig. 1.**
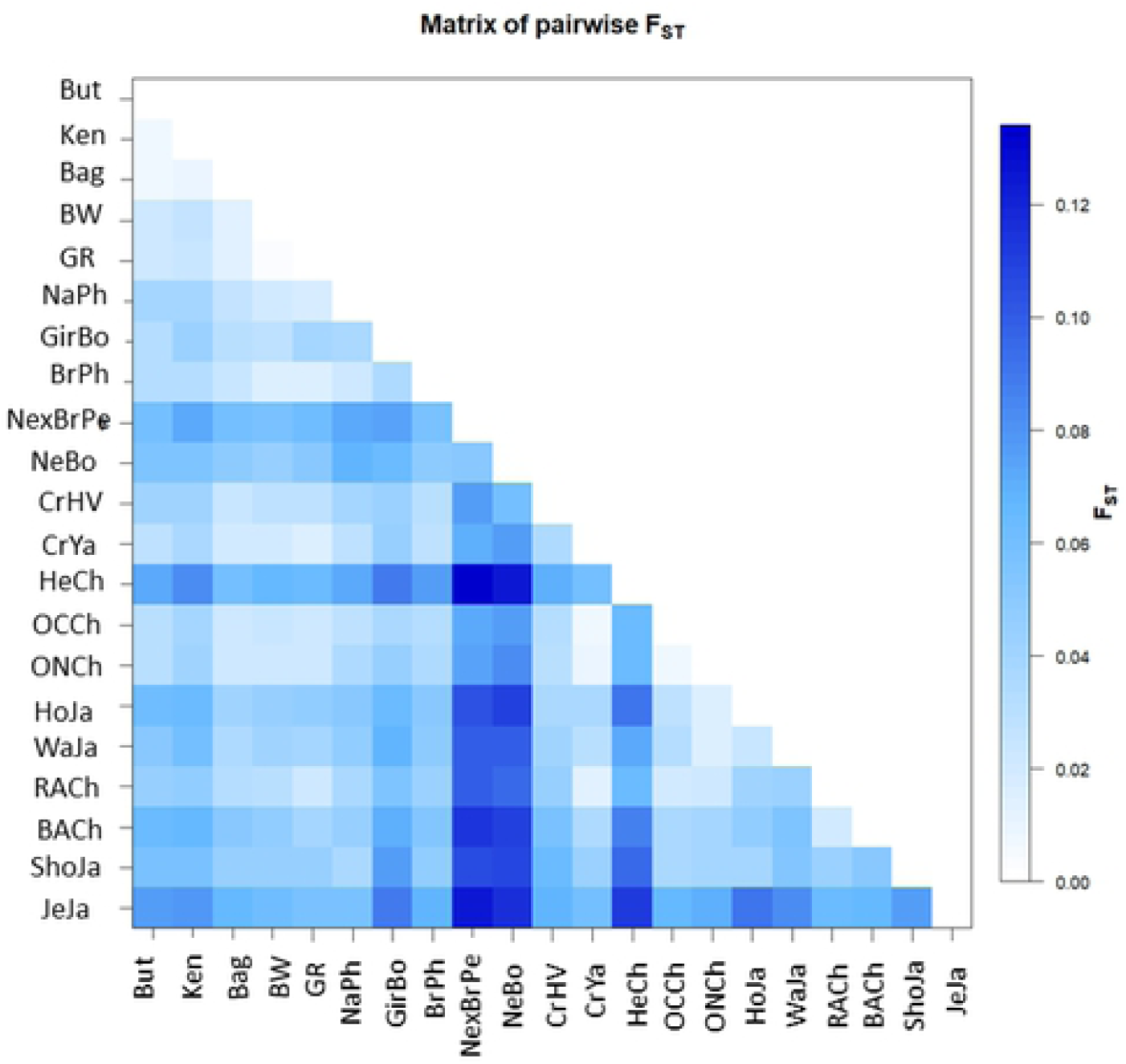
Alignment of the nucleotide (a) and the predicted amino acid (b) sequences of the β1 domain encoded by seven new *BoLA-DRB3* alleles (accession numbers, LC569725 for *BoLA-DRB3*\**004:02Sp*, LC569726 for *BoLA-DRB3*\**011:02Sp*, LC569729 for *BoLA-DRB3*\**018:01Sp*, LC569731 for *BoLA-DRB3*\**021:01sp*, LC569733 for *BoLA-DRB3*\**024:18Sp*, LC569735 for *BoLA-DRB3*\**027:05sp*, and LC569739 for *BoLA-DRB3*\**032:01sp*) derived from 225 Sudan native cattle cattle (113 animals of the Baggara native, 60 Butana, and 52 Kenana Sudan native cattle breeds). New alleles are indicated in bold. Numbering refers to amino acid positions in the mature protein. Nucleotide and amino acid residues identical to those encoded by the *BoLA-DRB3* cDNA clone NR-1 are indicated by dots (Aida *et al*., 1995). Missing data are indicated by dashes. Closer *BoLA-DRB3* alleles with new variants are also included in the figure. Id. = Nucleotide or amino acid identity in %.

The seven new variants were assigned allele names by IPD-MHC (https://www.ebi.ac.uk/ipd/mhc/group/BoLA; [34], namely, *BoLA-DRB3*\**004:02Sp*, which differs from *BoLA-DRB3*\**004:02* at two positions (141 and 197); *BoLA-DRB3*\**011:02Sp*, which differs from *BoLA-DRB3*\**011:02* at two positions (167 and 168); *BoLA-DRB3*\**018:01Sp*, which differs from *BoLA-DRB3*\**018:01* at one position (194); *BoLA-DRB3*\**021:01sp*, which differs from *BoLA-DRB3*\**021:01* at two positions (255 and 256); *BoLA-DRB3*\**024:18Sp*, which differs from *BoLA-DRB3*\**024:18* at one position (167); *BoLA-DRB3*\**027:05sp*, which differs from *BoLA-DRB3*\**027:05* at one position (219); and *BoLA-DRB3*\**032:01sp*, which differs from *BoLA-DRB3*\**032:01* at two positions (255 and 256). All seven new *BoLA-DRB3* allele variants shared about 89.7-92.6% and 80.52-85.71% nucleotide and amino acid similarity with the *BoLA-DRB3* cDNA clone NR1, respectively [35].

A Venn diagram was constructed using data obtained in this study and from previous reports [18,22–24,27]. Data were grouped in terms of the breed’s geographical origin as follows: native Sudanese (Baggara, Kenana, and Butana); Southeast Asian (Myanmar and Philippines native breeds); Zebu (Nellore, Gir, Brahman, and crossbreeds); European (Hereford, Black and Red Angus, Jersey, Shorthorn, Holstein, and crossbreeds); and American Creole cattle breeds (Overo negro, Overo Colorado). This analysis revealed that out of the 115 alleles identified in the five cattle groups, fourteen were unique to native Sudanese breeds (Fig. 2), four of which exhibited gene frequencies that were higher than 0.5%, representing about 26% of the 53 alleles detected in the native Sudanese cattle. In addition, two other variants were only present in native Sudanese and American Creole breeds, while six other alleles were only found in Sudanese cattle populations and American Creole or Southeast Asian native or Zebu breeds, or a combination of these groups. In addition, the BoLA-DRB3 NJ tree, including all the previously reported alleles and the seven new variants, showed that the variants detected in Sudanese cattle populations were interspersed among the various clusters (Fig. 3).

**Fig. 2.**
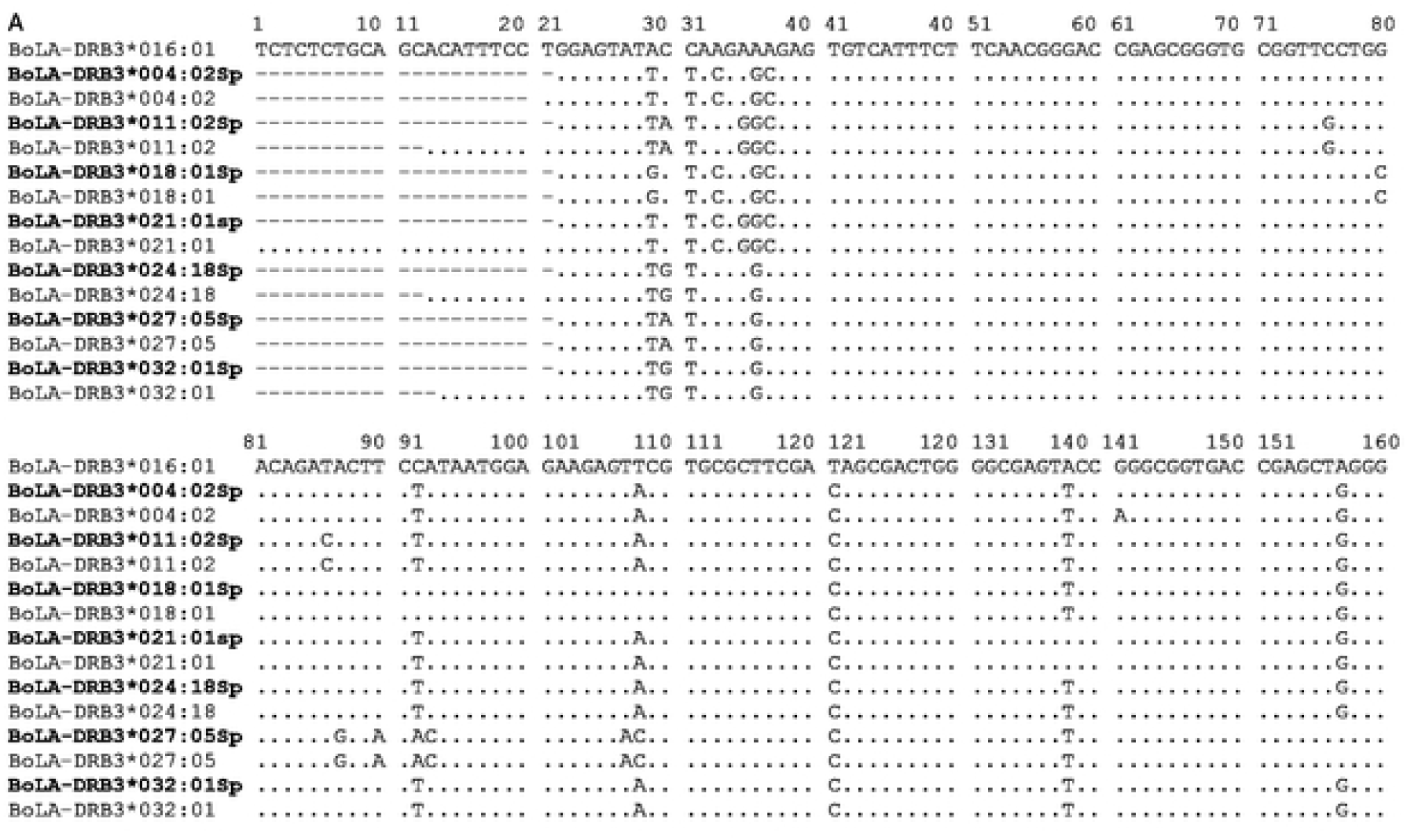
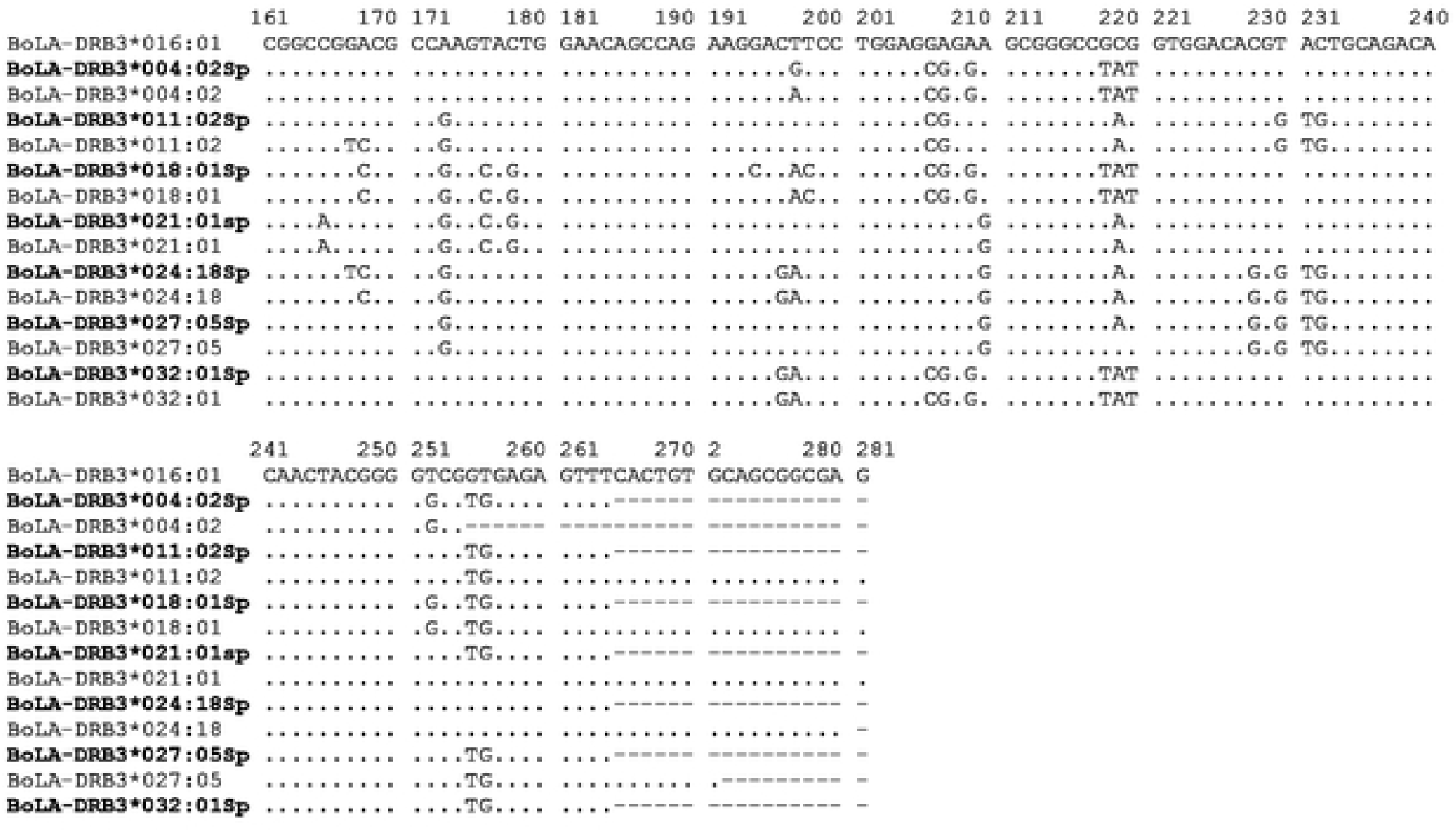
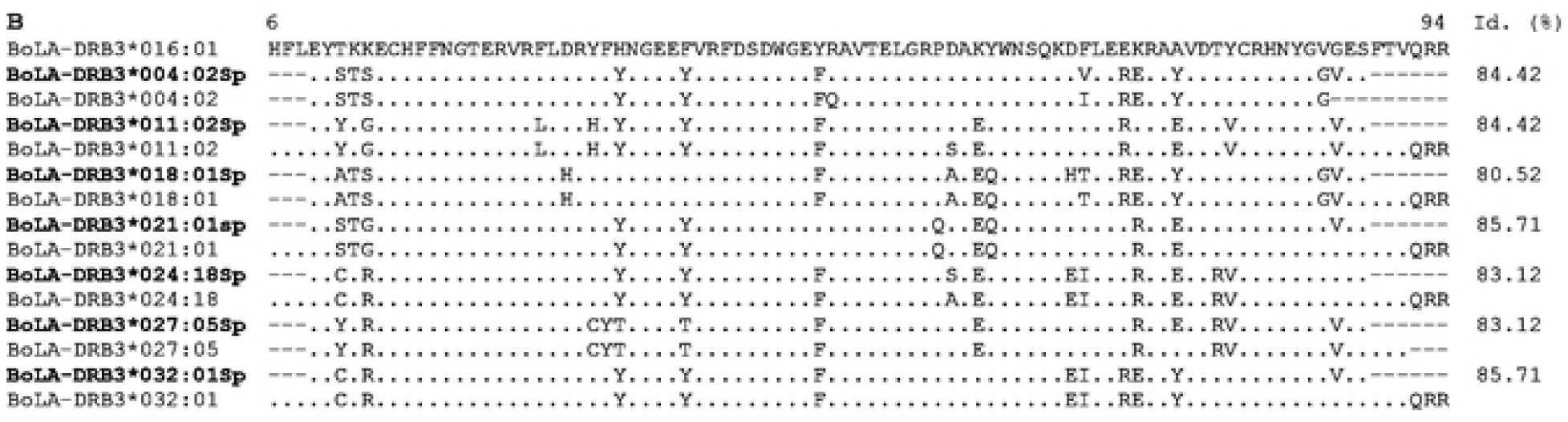
Venn plot of *BoLA-DRB3* alleles shared by Sudan native (Baggara, Kenana, and Butana); Southeast Asia (Myanmar and Philippine native breeds); Zebu (Nellore, Gir, Brahman, and crossbreeds); European (Hereford, Black and Red Angus, Jersey, Shorthorn, Holstein, overo negro, overo colorado, and crossbreeds); and American Creole (Yacumeño and Hartón del Valle) cattle breeds.

**Fig. 3.**
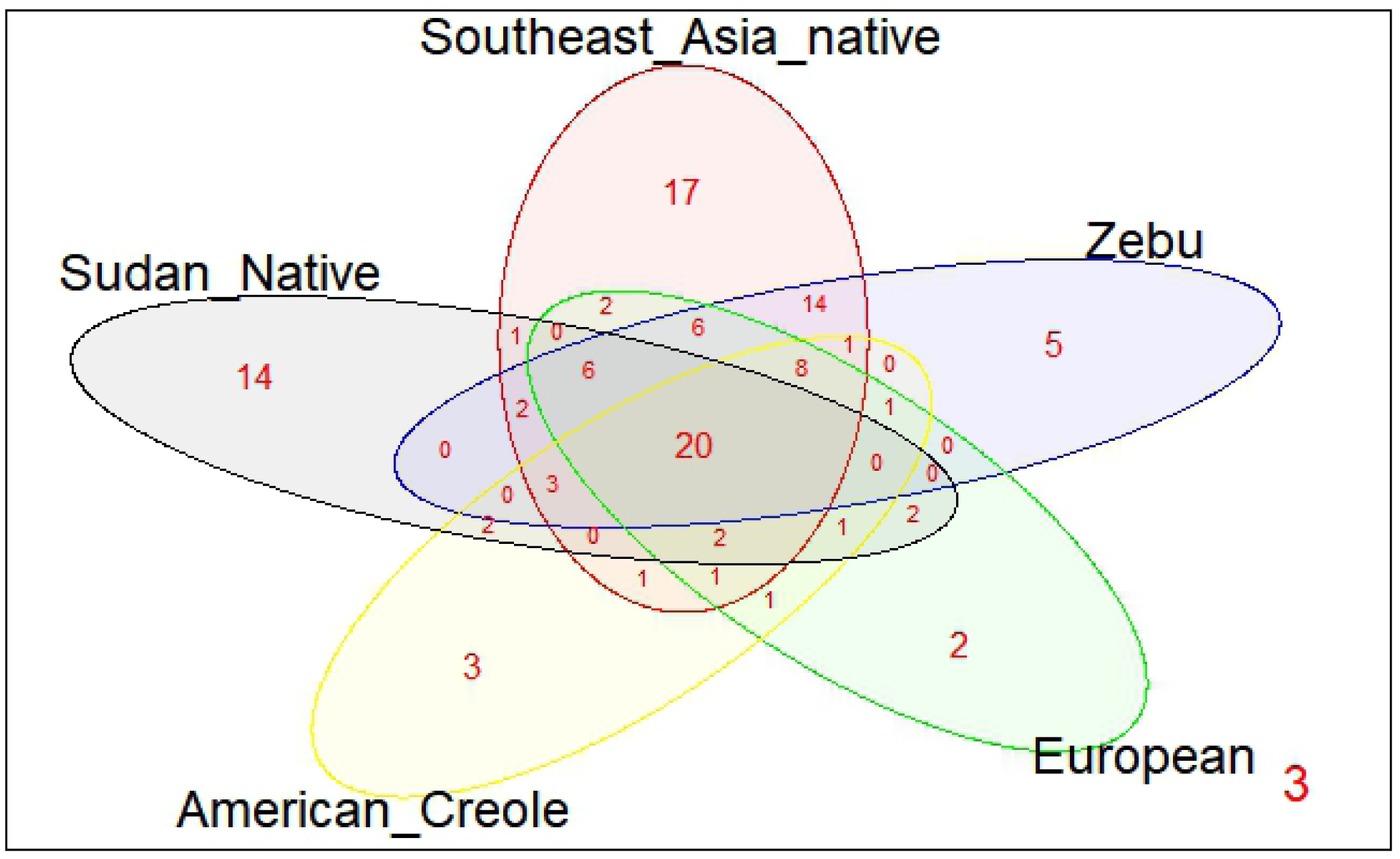
Graphic representation of calculated F_ST_ between population pairs using an R package pairFstMatrix.r. But = Butana, Ken = Kename, Bag = Baggara, BW = Pyer Sein. GR = Shwe Ni, NaPh = Philippine native, GirBo = Bolivian Gir, BrPh = Philippine Brahman, BrxNePe = Peruvian Brahman × Nellore crossbreed, NeBo = Bolivian Nellore, CrHV = Creole Hatón del Valle, CrYa = Creole Yacumeño, HeCh = Chilean Hereford, OCCh = Chilean Overo Colorado, ONCh = Chilean Overo Negro, HoJa = Japanese Holstein, WaJa = Japanese Black, BACh = Chilean Black Angus, RACh = Chilean Red Angus, ShJa =Japanese Shorthorn and JeJa = Japanese Jersey.

As shown in Fig. S1, the native Sudanese cattle breeds have an even gene frequency distribution, with a high number of alleles with low frequency. This was especially extreme in the Baggara breed. Only two, five and seven alleles appeared with frequencies of > 5% in the Baggarar, Kenana and Butana breeds, respectively. These common alleles accounted for a low proportion of the cumulative gene frequencies (12.83, 44.23 and 50.83% in the Baggara, Kenana and Butana breeds, respectively); four of which (*BoLA-DRB3*\**003:02:01*, \**021:01* \**022:01* and \**024:01*) were common in at least two out of the three Sudanese breeds (Table 1).

### Nucleotide and amino acid diversity in the *BoLA-DRB3* alleles found in native Sudanese cattle breeds

Genetic diversity at the DNA and amino acid levels was evaluated using four methods that compare the average amino acid and nucleotide substitutions for every pair of alleles within the breeds. The nucleotide (π) and the mean number of pairwise differences values within Sudanese native breeds were higher than 0.074 and 17.99, respectively (Table 2b). Comparison with results previously reported for other cattle breeds showed that these nucleotide diversity values all fall within the upper end of the range reported (π_range_ = 0.068-0.090; NPD_range_ = 16.31-20.96) when using PCR-SBT genotyping methods [18, 23–25, 27]. Regarding amino acid diversity, the average d_N_ and d_S_ substitutions in Sudanese cattle breeds were calculated across the entire *BoLA-DRB3* exon 2 and the antigen-binding site (ABS). As expected, the d_N_/d_S_ ratio was higher when only the ABS was analyzed (Table 2b). These values obtained in Sudanese cattle were similar to those estimated for other cattle breeds (d_N_/d_S total_ = 2.30 – 3.31; d_N_/d_S ABS_ = 2.63 – 4.11) [18, 23–25, 27].

### Gene diversity, Hardy-Weinberg Equilibrium (HWE), and neutrality testing of *BoLA-DRB3* variants found in Sudanese cattle breeds

Genetic diversity within the three Sudanese breeds was estimated using number of alleles (n_a_) and gene diversity (h_o_ and h_e_). Furthermore, we performed HWE and Slatkin’s neutrality tests on *BoLA-DRB3* to evaluate the possible effect of selection, inbreeding, and population structure on allelic diversity at this locus. The high n_a_ values and even gene frequencies observed in the Butana, Kenana and Baggara breeds resulted in h_e_ and h_o_ values higher than 0.93 (Table 2a). As expected, these indexes highlighted extremely high diversity values for Sudanese cattle populations, which is similar to the results reported for other bovine breeds which have been evaluated by PCR-SBT, and characteristic of *MHC class II DR* genes [18, 22–24, 27, 36]. Regarding the HWE test, the three Sudanese native populations were in equilibrium (Table 2a), similar to observations in half of the bovine breeds studied so far. It is widely accepted that the genetic diversity of *MHC class II* genes can be maintained by balancing selection. Thus, we performed a Slatkin’s exact neutrality test (Table 2a) to evaluate this phenomenon in the Sudanese cattle populations. The *BoLA-DRB3* gene frequency profile in Baggara cattle showed an even distribution (*p* = 0.016), consistent with the theoretical proportion expected under balancing selection pressures, as opposed to positive or neutral selection (*p* > 0.025). A similarly even *BoLA-DRB3* gene frequency was observed in other cattle breeds, including Japanese Black, Yacumeño Creole, Bolivian Gir, Pyer Sein and Shwe Ni. Conversely, *BoLA-DRB3* gene frequency distributions in Butana and Kenana cattle (*p* = 0.225 and *p =* 0.138, respectively) were more compatible with neutral selection, which is similar to the results obtained for the majority of the cattle breeds analyzed to date (Table 2a).

### BoLA-DRB3 genetic structure and levels of population differentiation in Sudanese cattle

The level of genetic differentiation among the three Sudanese breeds was studied through the *F*_ST_ index, and the results were then compared with the population structures common to the major groups (Taurine and Zebuine) and the bovine breeds already described in the literature. The average *F*_*ST*_ value showed a low level of genetic differentiation (*F*_*ST*_ = 0.0076) among native Sudanese breeds, ranging between 0.007 and 0.009 (Table S3). This average value is higher than those estimated in Myanmar native breeds (*F*_*ST*_ = 0.003), and slightly lower than those reported for Holstein populations from different countries (*F*_*ST*_ = 0.009) [24,29] (Figure 4 and Table S2). When breeds were grouped in terms of the breed’s geographical origin, as was done in the Venn diagram, the genetic variance among breed groups and among populations within groups accounted for 1.18% and 3.71% of the total genetic variance. Table S3 summarizes the genetic distance, measured by *F*_ST_, between native Sudanese breeds and other Taurine and Zebu breeds for BoLA-DRB3, showing that native Sudanese cattle diverge from other breeds with *F*_*ST*_ values between 0.014 and 0.082.

**Fig. 4.**
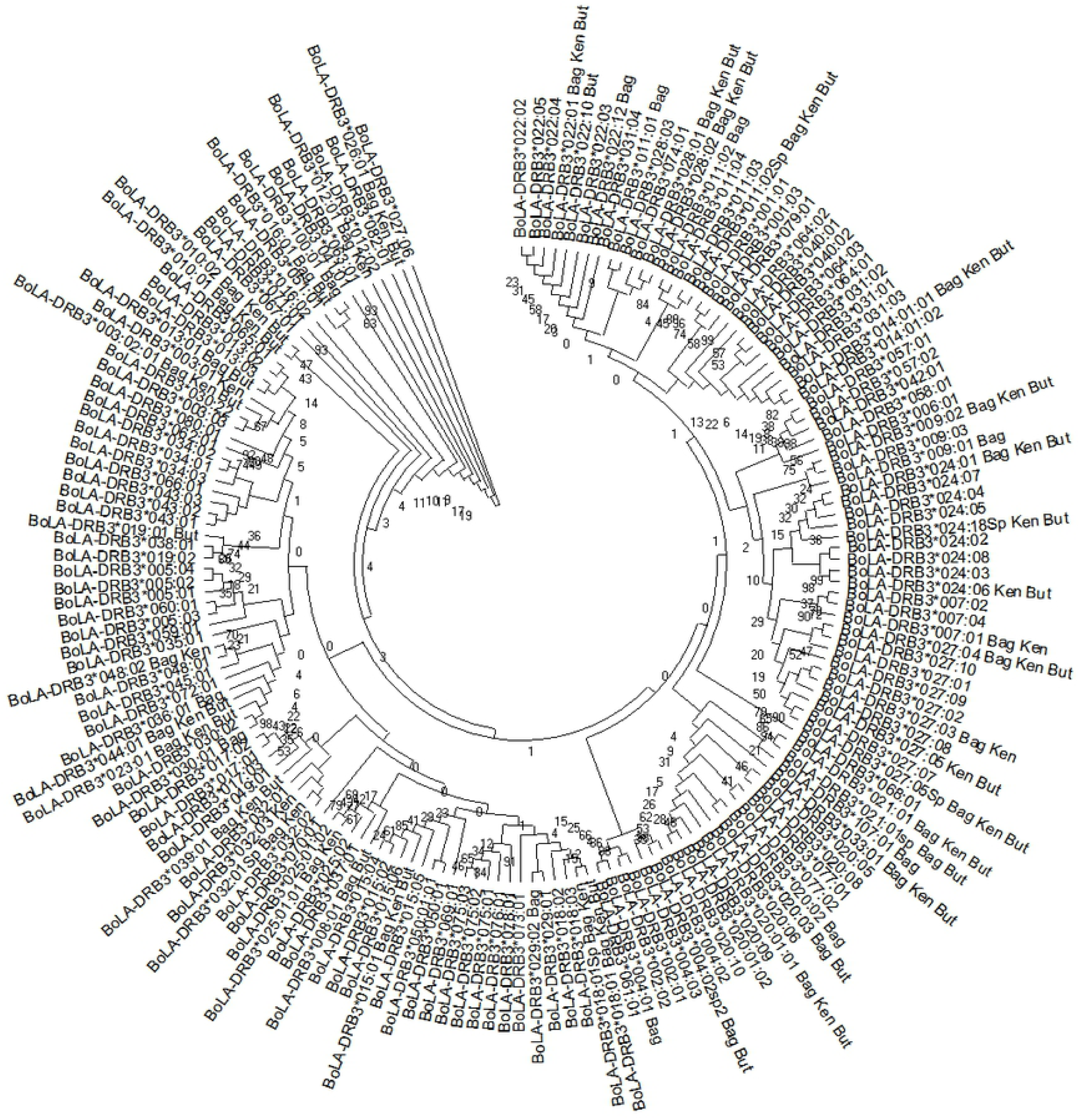
Neighbor-joining (NJ) tree constructed from the 270 bp nucleotide sequence that includes the β1 domain encoded by all reported *BoLA-DRB3* alleles and the seven new ones (*BoLA-DRB3*\**004:02Sp2, BoLA-DRB3*\**011:02Sp, BoLA-DRB3*\**018:01Sp, BoLA-DRB3*\**021:01sp, BoLA-DRB3*\**024:18Sp, BoLA-DRB3*\**027:05sp*, and *BoLA-DRB3*\**032:01sp*). Numbers are bootstrap percentages that support each node. Bootstrapping was carried up with 1000 replicates to access the reliability of individual branches. Bag = Baggara, But = Butana, Ken = Kenana.

When the five sample sites of native Sudanese breeds were compared (two sites of Kenana cattle were very close and assumed as one), the average *F*_ST_ value was 0.0074 (*p* = 0.164), while the pairwise *F*_ST_ ranged from 0.0002 (p = 0.450) between both Baggara populations and 0.0118 (p < 0.0001) between Baggara Daiwani and Butana Bu_Qadarif. Significant differences were observed in nine out of the ten native population comparisons (*p* < 0.05; Table S4). Similar genetic distance values were observed among Holstein populations from different countries and between native breeds of Myanmar [24, 29].

### Genetic differentiation of *BoLA-DRB3* alleles in native Sudanese cattle breeds: Comparison with Zebu and Taurine breeds

First, BoLA-DRB3 allele frequencies from Sudanese cattle populations and for each breed included in the dataset were used to generate Nei’s DA and DS genetic distance matrices. Then, dendrograms were constructed from these distance matrices using UPGMA and NJ algorithms. All trees revealed congruent topologies, which were consistent with the historical and geographical origin of the breeds analyzed. As expected, these trees revealed two main clusters which included the Taurine and Zebuine breeds, respectively (Fig. 5a). It is noteworthy that Sudanese breeds were located in a sub-cluster within the Zebuine cluster, with the two dairy breeds located in the east of the country, Butana and Kenana, more related to each other with respect to the Baggara beef breed in the west. These results reveal that Sudanese cattle breeds have a particular diversity in the BoLA-DRB3 gene, as a consequence of its gene frequency profile and the presence of a high number of private alleles.

**Fig 5.**
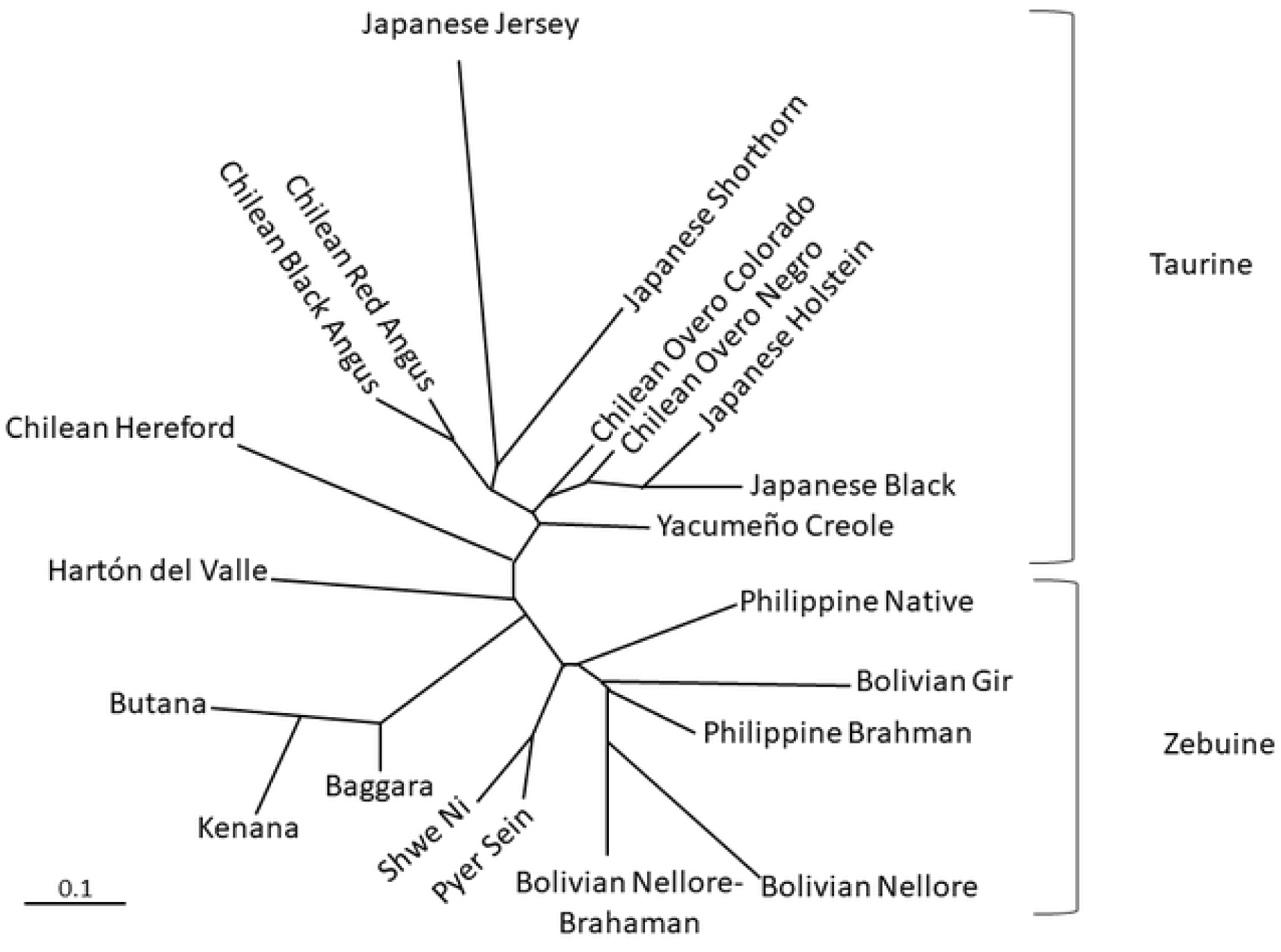
Neighbor-joining dendrogram constructed from a matrix of D_A_ genetic distances. (b) principal component analysis of allele frequencies from the BoLA-DRB3 gene in 22 breeds. But = Butana, Ken = Kename, Bag = Baggara, BW = Pyer Sein. GR = Shwe Ni, NaPh = Philippine native, GirBo = Bolivian Gir, BrPh = Philippine Brahman, BrxNePe = Peruvian Brahman × Nellore crossbreed, NeBo = Bolivian Nellore, CrHV = Creole Hatón del Valle, CrYa = Creole Yacumeño, Creole Highland, HeCh = Chilean Hereford, OCCh = Chilean Overo Colorado, ONCh = Chilean Overo Negro, HoJa = Japanese Holstein, WaJa = Japanese Black, BACh = Chilean Black Angus, RACh = Chilean Red Angus, ShJa =Japanese Shorthorn and JeJa = Japanese Jersey.

The results of the PCA based on the *BoLA-DRB3* allele frequencies from 22 breeds showed that the first three components accounted for 47.30% of the data variability. The first PC accounted for 24.31% of the total variance and, as shown in a previous study [25], clearly exhibited a differentiation pattern between the Zebu (negative values) and Taurine (positive values) breeds, while native breeds from Southeast Asia and Sudan were located in an intermediate position near the axis origin of the plot (Fig. 5b). The first PC was primarily determined by differences in the frequency of the same alleles, such as BoLA-DRB3*022:01, *028:01, *036:01, *031:01, *030:01, and *057:02 with the higher negative axis 1 values, while, the alleles BoLA-DRB3*001:01, *002:01, *007:01, *008:01, *010:01, *011:01, *012:01, *015:01 *016:01, *018:01 had the higher positive values for this axis. The second PC explained 11.98% of the total variation and showed a gradient among Taurine breeds, with Chilean Hereford (positive values) and Japanese Jersey (negative values) located at opposite ends. Furthermore, this component discriminated between native Sudanese and native Southeast Asian cattle breeds. Finally, the third PC accounted for 11.01% of the variance and allowed for the differentiation of Chilean Hereford, and Japanese Jersey and Japanese Holstein cattle from other Taurine breeds. In summary, the native Sudanese cattle breeds were located within a narrow cloud in an intermediate position between the Zebu and Taurine breeds and close to other Southeast Asian breeds, in agreement with the composite origin of these native breeds, which is also supported by the presence of African and Zebu unique *BoLA-DRB3* alleles within these populations. These PCA results agree with the overall clustering observed after NJ or UPGMA tree construction.

The BoLA class II molecule binds peptides derived from antigens via five antigen binding pockets named pocket 1, pocket 4, pocket 6, pocket 7 and pocket 9 [20]. To assess whether observed differences in allelic frequency is reflected within amino acid motifs in each pocket, we analyzed frequency of the protein pockets implicated in the antigen-binding function of the MHC complex by PCA. As shown in Figure S2a-e, the three native breeds of Sudan are located in a closed cloud in the five PCAs made based on the frequency of the pockets, although varying their relative position with other breeds and breed groups, and in some cases the spatial distribution did not exhibit a clear relationship with the geographical or historical origin of the breeds. However, pockets 4 and 9 are the ones that best differentiate these native breeds from the rest. Regarding pocket 4, the native breeds of Sudan are located in a narrow cloud located at the end of axis 2, and their position is mainly explained by the GFDEREY, RFDERFV and GLDRKEV motifs. The position of the native Sudanese breeds in pocket 9 was the result of positive PC1 and PC2 values for the presence of amino acid motifs EYD and EFA.

Finally, PCA was performed at the Sudanese population level to evaluate the degree of genetic structure among the sampling sites (Baggara Daiwani, Baggara Nyakawi, Kenana, Butana Bu Atbara and Butana Bu Qadarif). This analysis showed that the first three components accounted for 90.95% of the data variability. The first PC accounted for 30.65% of the total variance and clearly exhibited a differentiation pattern between the Baggara population (negative values) and the Butana Bu Qadarif (positive values) population, while Kenana, Butana Bu Atbara were located in intermediate positions (Fig. 6). These results agree with the geographical distribution of the studied population. The second and third PCs explained 30.66% and 25.24% of the total variation and allowed for the differentiation of the Butana Bu_Qadarif and Kenana populations, respectively.

**Fig 6.**
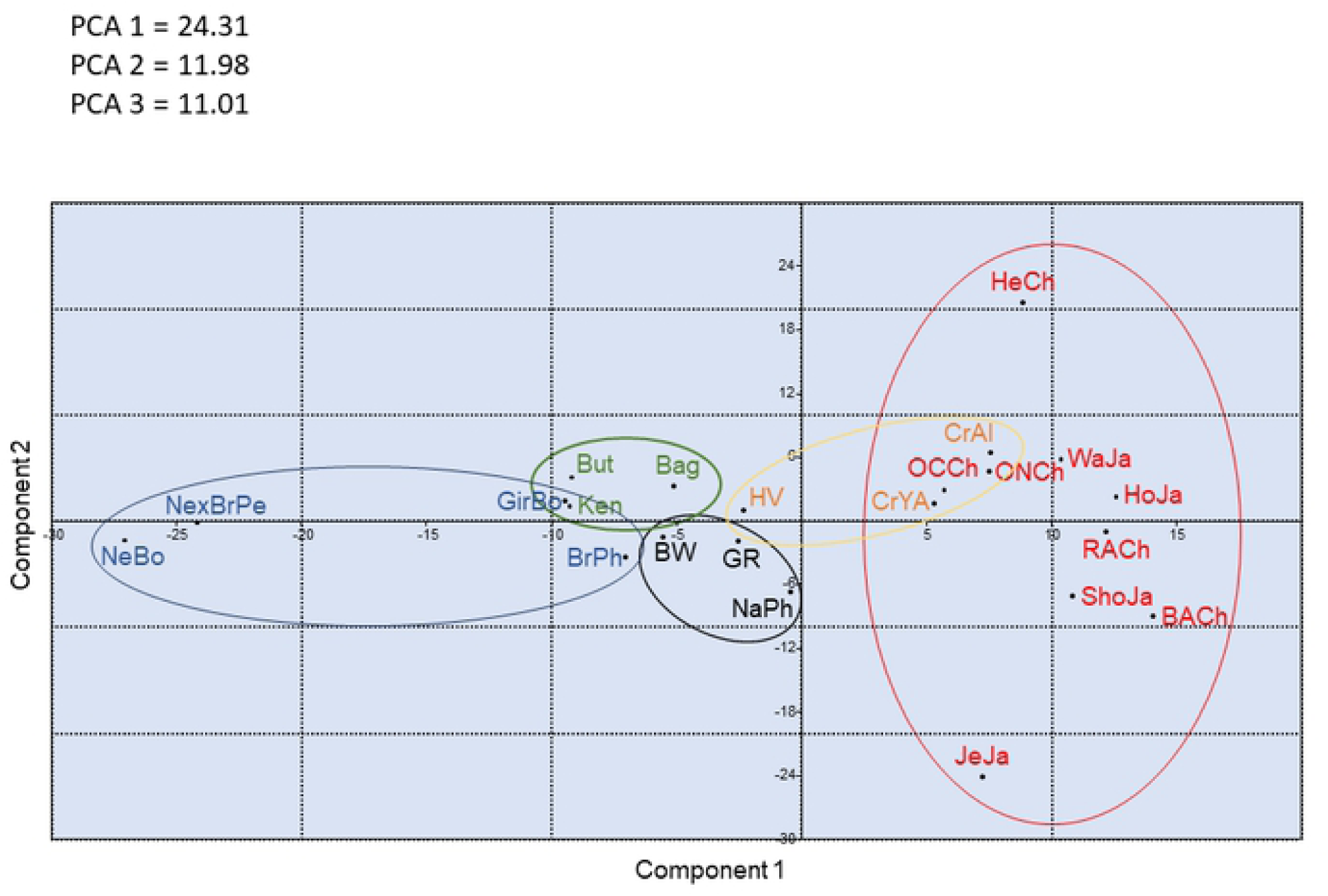
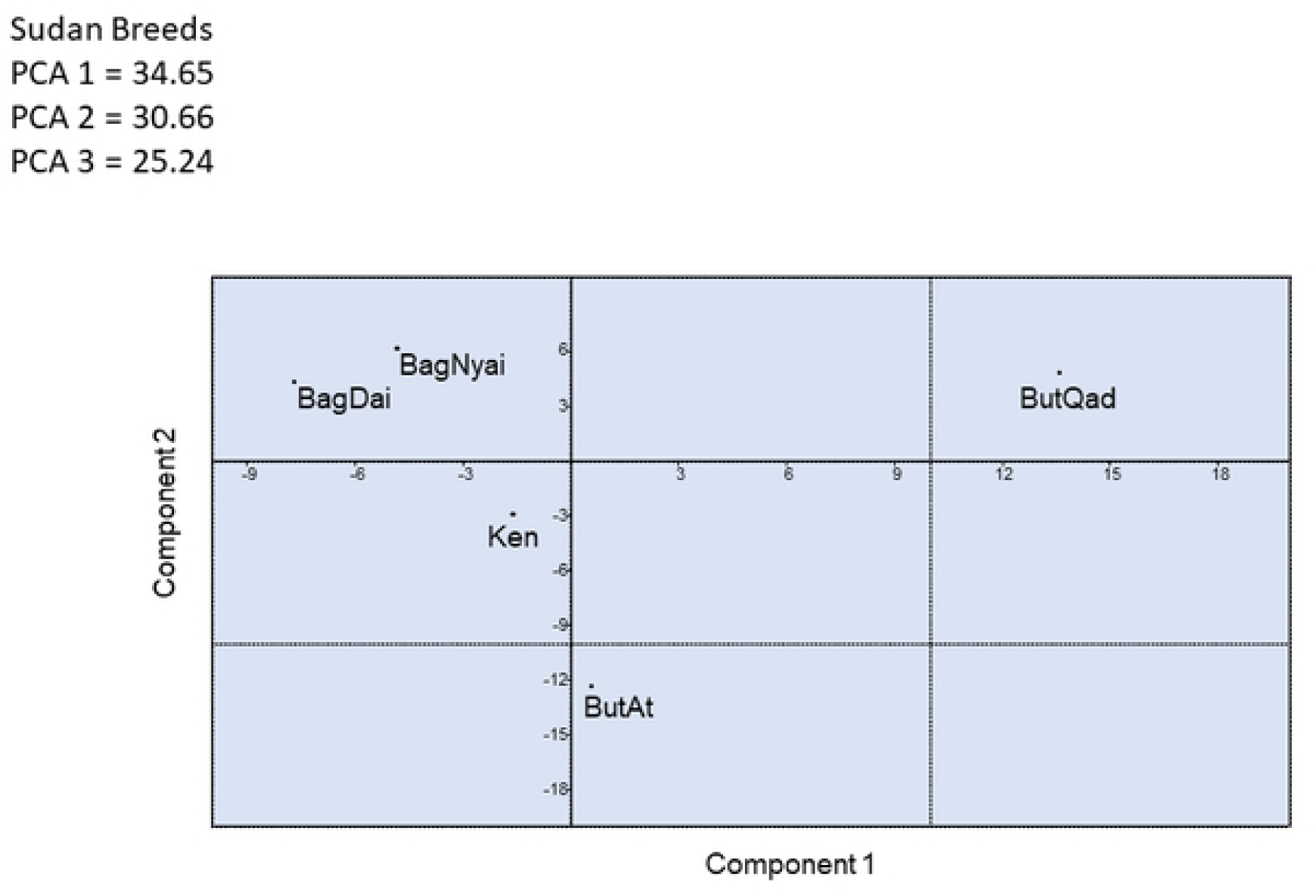
Principal components analysis of allele frequencies from the BoLA-DRB3 gene in five Sudan native samples sites (BagDai = Baggara Daiwani, BagNyai = Baggara Nyakawi, Ken = Kenana, ButAt = Butana Bu Atbara, and ButQad = Butana Bu Qadarif).

## Discussion

Since the first pioneering studies based on serotype analysis, a number of striking differences between the BoLA profiles of African and European cattle have been reported due to difference in the antigen’s frequency of occurrence and the presence of unique antigens in African cattle [37]. Over the next decades, several private alleles were identified in Taurine, Zebu and Taurindicus native African breeds, like N’Dama, Boran, and Sanga [31–33]. However, in the present study, we carried out the first genetic characterization of the *BoLA-DRB3* gene at population level in native Sudanese breeds using PCR-SBT. This analysis allowed us to detect 53 alleles, including seven new variants. The high number of private alleles agrees with data obtained by Kim et al. [15], who analyzed the BoLA region in depth using a genome-wide sequencing approach, identifying six major African BoLA haplotype blocks.

The wild cattle or aurochs (*Bos primigenius*), the ancestor of domestic cattle, inhabited a large geographical area throughout Eurasia and North Africa. According to the trans-specific theory of MHC alleles [38], it is expected that the extremely high genetic variability present in the BoLA-DRB3 gene (330 alleles have been reported in the IPD database, access date 07/06/20) was present in the wide geographical distribution of the aurochs. On the basis of archeological and genetic studies, it has been proposed that bovines were domesticated in five geographical sites (domestication centers), one located in the Near East, one in North Africa, two in India and one in Northeast Asia [39–43]. From the domestication centers, domesticated bovines spread throughout the continents, following the same migration routes as human populations. Each of these domestication centers would have retained only a fraction of the total diversity as a result of bottleneck and genetic drift effects [44]. This is clearly seen in the distribution of mitochondrial haplogroups among cattle breeds, where T, T2 and T3 haplogroups are mainly retained in the Near East domestication center, T1 is predominant in Africa cattle breeds, T4 is detected in Northeast Asia, and I1 and I2 in India [4, 39–43]. In addition, it was postulated that the remaining haplogroups, Q and P, were introgressed through the crossbreeding between local wild aurochs and domestic cattle. In Africa, Taurine cattle would have been domesticated in the north-east of the continent and from there they would have dispersed south and west. Then, Zebu cattle were introduced and indicus genes were introgressed into native populations through absorbent crosses [45]. Currently, an east-west gradient of Zebu influence in African native genes is observed.

Subsequent dispersal and crossbreed processes (migration and gene introgression) and natural and artificial selection would have shaped the BoLA-DRB3 diversity within the current bovine populations. Accordingly, the *BoLA-DRB3* alleles detected in the Sudanese cattle were interspersed distributions along the allele NJ tree instead of grouped in specific clusters of the dendrogram, which is consistent with the ancient origin of the BoLA-DRB3 alleles. Similar results have been reported in other native cattle breeds from different geographical regions [27, 29].

In order to understand how the successive processes of domestication, migration and adaptation shaped the distribution of the *BoLA-DRB3* alleles within native Sudanese breeds, a Venn diagram was constructed. This analysis demonstrated that 14 *BoLA-DRB3* alleles were only detected in Sudanese cattle populations, at least for the breeds included in this analysis. Seven of these alleles corresponded to new variants described in this study (*BoLA-DRB3*\**004:02Sp2, BoLA-DRB3*\**011:02Sp, BoLA-DRB3*\**018:01Sp, BoLA-DRB3*\**021:01sp, BoLA-DRB3*\**024:18Sp, BoLA-DRB3*\**027:05sp*, and *BoLA-DRB3*\**032:01sp*). Furthermore, a review of the IPD–MHC database showed that this group of Sudanese private alleles included seven other variants previously detected in African breeds (BoLA-DRB3*022:10; BoLA-DRB3*022:12; BoLA-DRB3*024:01; BoLA-DRB3*024:06; BoLA-DRB3*032:03; BoLA-DRB3*100:01; BoLA-DRB3*107:01; Table S4).

Two BoLA-DRB3 alleles, that were only previously reported in Creole cattle breeds [27], were identified in native Sudanese breeds. Studies based on mitochondrial DNA and Y chromosome haplotypes have revealed an African component in the germplasm of the American creole bovine breeds. Two origins have been proposed for this African component: through the native Iberian cattle that originated the Creole cattle and/or a direct introgression from mainland Africa following the slave trade routes [46]. The Iberian theory is unlikely as the BoLA-DRB3*011:02 and BoLA-DRB3*029:02 alleles have not been detected in the Spanish Morucha breed, which is the only autochthonous Iberian breed in which the genetic diversity of the BoLA-DRB3 gene has been studied [28]. In summary, 16 possible African putative alleles were identified in the native bovine populations of Sudan, totaling 20.22% of the gene frequency.

On the other hand, a group of alleles is shared between the Sudanese breeds and the Zebu, Southeast Asian and/or Creole American breed groups (BoLA-DRB3*003:02:01, BoLA-DRB3*005:02, BoLA-DRB3*020:03, BoLA-DRB3*023:01, BoLA-DRB3*025:01:01, BoLA-DRB3*027:04, BoLA-DRB3*030:01 and BoLA-DRB3*039:01), but is absent in the European breeds. It is worth noting that these alleles were first identified in cattle breeds such as Boran, Ethiopian Arsi, N’Dama and Brahman [31–33, 47] (Table S5). The introgression of these variants could be a consequence of the successive waves of introduction of Zebu cattle into the African continent [45]. These alleles account for an additional 15.33% of the gene frequencies. The remaining alleles have a worldwide geographical distribution, thus 20 variants have been detected in all the breed groups included in the Venn diagram. Further studies on the genetic diversity of the BoLA-DRB3 gene in other African bovine populations will surely reveal a greater allelic repertoire in these important animal genetic resources. In this sense, a review of the IPD–MHC database showed that some variants which had been previously detected in African Zebu breeds (*BoLA-DRB3*\**038:01* in Ethiopian Arsi and *BoLA-DRB3*\**013:03* in Sudanese Baggara) were not identified in the present studied sample.

The current repertoire of alleles of the BoLA-DRB3 gene in the native cattle of Sudan would not only have been molded by stochastic forces, such as the formation of the founder group, gene drift and recent or historical gene introgression as described above, but also by processes of natural and artificial selection. In Sudan, as in other African regions, cattle are subjected to strong environmental pressures, such as tropical diseases, heat stress, drought and poor nutritional and forage deficits. Furthermore, animals are affected by diverse infectious diseases, including parasites (e.g., ticks, theileriosis, babesiosis, anaplasmosis, trypanosomosis; [48–51], bacteria (e.g., Hemorrhagic septicemia, Anthrax, tuberculosis, brucellosis, Thrombotic meningoencephalitis; [52–56] and viruses (e.g., foot and mouth disease, lumpy skin disease, Pox virus, bovine viral diarrheal diseases complex; [57–59]. For this reason, it is to be expected that native Sudanese cattle will be under strong selection pressure, which would contribute to maintaining and shaping the genetic diversity of the BoLA-DRB3 gene. In this sense, a wide repertoire of alleles allows the population to identify and respond to a greater range of antigens. Furthermore, heterozygous animals would trigger an immune response to a greater variety of antigens. For these reasons, it has been proposed that this allelic diversity is maintained by balancing or over-dominant selection [25,60,61]. Different indices at the population, nucleotide and amino acid levels showed high levels of genetic diversity in the bovine breeds of Sudan for the BoLA-DRB3 gene. This is clearly reflected in the presence of a homogeneous distribution of gene frequencies (a high number of alleles with low frequencies). This is particularly extreme in the Baggara breed in which Slatkin’s neutrality test evidenced that the *BoLA-DRB3* gene frequency profile showed an even distribution consistent with the theoretical proportion expected under balancing selection pressures. Similar results have been reported for other cattle breeds, including Japanese Black, Yacumeño Creole, Bolivian Gir, Pyer Sein and Shwe Ni [25, 27, 29].

In contrast, the HWE analysis did not show a significant increase in heterozygous individuals in the Sudanese cattle populations studied, which could be a consequence of the effect of over-dominant selection [62]. As discussed in previous works [27], this effect has been observed only in some of the breeds studied so far and the most common explanation for the absence of heterozygote excess in the studied bovine breeds is the low magnitude of the overdominance selection coefficient at MHC loci (probably lower than 0.02; [63]). Such weak selection would only be enough to increase the number of heterozygotes in large populations and in the absence of high rates of stochastic forces (population bottlenecks, genetic drift, and inbreeding). For this reason, and because the HWE method may suffer from low resolving power, such effects were not observed.

The repertoire of alleles of the BoLA-DRB3 gene present in the native cattle of Sudan allows these breeds to be clearly differentiated from the rest, forming a cluster in the NJ and UPGMA trees and a narrow cloud in the PCA. This pattern is confirmed when PCAs are performed based on the pocket 4 and 9 gene frequencies. Noteworthy is that it has been proposed that pocket 4 plays an important role in the binding of peptides due to this pocket being located in the center of the PBC [64,65]. In addition, it has been reported in cattle that immune responses against vaccine and disease resistance is significantly related to differences in the pocket 4 motif [46,47]. In this sense, a particular amino acid (e.g., amino acid R in position 70) or amino acid motifs (e.g., ER at 70 and 71 sites; EIAY motif at positions 66–67–74–78, and the deletion of the amino acid 65), in sites that affect the conformation of pocket 4, have been associated with immune response or resistance to infectious diseases, such as mastitis, persistent lymphocytosis, dermatophilosis, and tick-born diseases [19,64-69]. Many of these diseases, as well as others mentioned above, are present in Sudan and could have contributed to shaping the current repertoire of BoLA-DRB3 alleles present in native Sudanese cattle. However, these results were obtained in breeds that have different genetic backgrounds and that are raised in different environments and production systems, so further association studies are necessary to determine the effect (resistance or susceptibility) of the alleles present in the native cattle breeds of Sudan against different infectious diseases.

## Conclusions and future prospects

To the best of our knowledge, this is the first study to document in detail the genetic diversity and ancestral origin (Taurine vs Zebu) of BoLA-DRB3 alleles in cattle not only in Sudan but in the entire African continent. In addition to the good clustering of cattle based on ancestral origin and phylogeography, we identify seven novel alleles in the three native Sudanese cattle breeds. Two evolutionary forces contribute to the preservation and shaping of the genetic diversity of the BoLA-DRB3 gene in native Sudanese cattle; balancing selection for the Baggara breed and positive or neutral selection for the Butana and Kenana breeds. The results demonstrate that the background variation between two cattle groups, Taurine and Zebu, is primarily due to events of origin, selection, and adaptation, which explains the variations found in the diversity of the BoLA-DRB3 genes, not only between the two major groups but also with the Zebu cattle group, as documented in this study. This variation may explain how these cattle from Sudan are well adapted and resistant to various diseases. We presume that this genetic information provides a basis for better design of suitable breeding schemes. Further studies are highly encouraged to link the newly identified variants and their association with specific common disease(s).

This study has some limitations. Inclusion of more breeds from Sudan may have given a more comprehensive picture. In addition, the unavailability of data for African cattle against which to make comparisons is a shortcoming to this study.

## Materials and methods

### Sampled populations and genomic DNA extraction

The ODK system was used to record the sampling information; breed name, sex, estimated age, sampling location GPS coordinates, photo of the animal and owner’s information. Three cattle breeds were sampled: 1) Butana breed: collected from Atbara Butana Station and surrounding villages and from El-Gadarif city and Butana plain; coat color is red, brown and very rarely white, eyes are surrounded by dark black color as well as a dark black color at the ankle. 2) Kenana breed: samples were collected from Rabak city and surrounding villages and from UmBanein Kenana Station; it is characterized by a white color, somewhere greyish and a dark black color at the neck and face, dairy cattle structure. 3) Baggara breed: two populations of western Baggara breed were sampled; Nyalawi: white or white with some black splashes, big in size, beef cattle structure sampled from calves from Nyala city, South Darfur. Daeinawi from Ed daein city; Coat color red, beef cattle structure, smaller in size, as compared with Nyalawi, this population is characterized by a deep dark red coat color and black color along the neck, the lateral side of the head, the hind quarters and shoulder sides (Fig. S3).

A total of 225 animals were sampled (Baggara (N = 113), Butana (N = 60) and Kenana (N = 52) native breeds (Table S1 and Fig. S3)). Seven milliliters of venous blood were collected in EDTA-containing vacutainer tubes. Genomic DNA was extracted using DNeasy® Blood and Tissue Kit, (Qiagen, Germany), following the manufacturer’s instructions.

### PCR amplification and sequencing

Exon 2 of the BoLA-DRB3 was amplified by PCR as described by Takeshima *et al*. [21]. Using DRB3FRW 5-CGCTCCTGTGA(C/T)CAGATCTATCC-3 and DRB3REV 5-CACCCCCGCGCTCACC-3, PCR reactions were performed in a 25μl-reaction mixture containing 12.5μl of 2 × Gflex PCR Buffer (Mg2+, dNTP plus) (TaKaRa Bio Inc., Shiga, Japan), 0.5μl of Tks

Gflex DNA polymerase (1.25 units/μl) (TaKaRa Bio Inc.), 200nM of each primer, and 1.0μl of template. The reaction conditions consisted of an initial denaturation step at 95°C for 3min, followed by 35 cycles of 95°C for 1min, 58°C for 30s and 68°C for 90s and a final extension step at 68°C for 5min. PCR products were purified using a NucleoSpin Gel and PCR Clean Up Kit (Takara Bio Inc.). Cycle sequencing reactions were performed directly using the two PCR primers using the BigDye Terminator version 3.1 Cycle Sequencing Kit (Applied Biosystems, Foster City, CA, USA) and analyzed on an ABI Prism 3130 x genetic analyzer (Applied Biosystems) according to the manufacturers’ instructions.

### Sequence data analysis

Prior to analysis, all the chromatograms were visualized and sequence fragments were edited manually using ATGC software version 9.1 (GENETYX Corporation, Tokyo, Japan) correcting base calling errors. Multiple sequence alignments were performed using the MUSCLE algorithm implemented in MEGA X [70], and were subsequently joined to reconstruct a fragment of 280bp spanning the entire exon 2. The sequences obtained for the seven new alleles in the present study were submitted to the DNA Data Bank of Japan (http://www.ddbj.nig.ac.jp) under accession numbers LC569724-LC569739.

### BoLA-DRB3 allele genotyping

For typing BoLA-DRB3 genotypes, we constructed a new allele database using the IPD-MHC database. FASTA file, MHC_nuc.txt, was downloaded from the IPD-MHC Download page (https://www.ebi.ac.uk/ipd/mhc/download/) and BoLA-DRB3 allele sequences were extracted. This FASTA file was used as the initial BoLA-DRB3 database. Both sides of the sequenced raw data (ab1 files) from each cow were subjected to Assign 400ATF ver. 1.0.2.45 software (Conexio Genomics, Fremantle, Australia), with constructed the BoLA-DRB3 database, and performed genotyping. When we found a clear mismatch from several samples, we assigned these samples containing new alleles and revised the BoLA-DRB3 database containing new allele sequences. If the sample could not genotype using these criteria, we discarded the sample result from this analysis.

### Statistical analyses

#### Genetic diversity at allele level

Allele frequencies and the number of alleles (n_a_) were obtained by direct counting. The distribution of alleles across breeds was analyzed by a Venn plot created using the R package ‘VennDiagram’ (http://cran.r-project.org/. The observed (h_o_) and unbiased expected (h_e_) heterozygosity of the *BoLA-DRB3* locus were estimated according to Nei [71] using the Arlequin 3.5 software for population genetic analyses [72]. *F*_IS_ statistics [73] for each breed were calculated using the exact test included in Genepop 4.7 software [74] to evaluate deviation from Hardy-Weinberg equilibrium (HWE). The Ewens–Watterson–Slatkin exact test of neutrality was carried out using the method described by Slatkin [75] and implemented in the Arlequin 3.5 program.

#### Breed genetic structure

Genetic structure and genetic differentiation within Sudanese cattle breeds and among bovine breeds were assessed using Wright’s *F*_ST_ statistics [73]. This parameter was estimated using Arlequin 3.5 and Genepop 4.7 software. The *F*_ST_ values were represented graphically using the pairFstMatrix.r function implemented in the R statistical environment.

#### Genetic relationship between breeds

To condense the genetic variation at the *BoLA-DRB3* locus, allele frequencies were used to perform a Principal Component Analysis (PCA) according to the Cavalli-Sforza and co-authors [76] method, implemented in Past software [77]. Nei’s standard genetic distances Ds [78] and D_A_ [79] were calculated from allele frequencies and were used to perform cluster analysis using the unweighted pair-group method with arithmetic mean (UPGMA) [80] and the neighbor-joining (NJ) algorithm [81]. Confidence intervals for the groupings were estimated by bootstrap resampling of the data using 1000 replicates. Genetic distances and trees were computed using the Populations 1.2.28 software [82]. The trees were then visualized using TreeView [83].

#### Genetic diversity at sequence level

Nucleotide diversity (π) and pairwise differences in nucleotide substitutions between alleles within each breed were calculated using Arlequin 3.5. The mean number of nonsynonymous (d_N_), and synonymous (d_S_) nucleotide substitutions per site from averaging over all sequence pairs were estimated within each group using the Nei-Gojobori model [84] and Jukes– Cantor’s formula implemented in the software MEGA X (70). The *BoLA-DRB3* allele tree was constructed from a distance matrix that was based on the NJ method using the MEGA X software. To test the significance of the branches, 1000 bootstrap replicate calculations were performed.

## Ethical approval and consent to participate

This study was approved by the Faculty of Veterinary Medicine Research Board, University of Khartoum, Sudan (Approval no. 1-2017). Consent of the animal owners was sought before blood samples were collected.

## Authors’ contributions

BS and JMM conceived the initial study, BS designed all the described experiments. B.S, S-NT GG, analyzed the data and drafted the manuscript. M-KA, helped in drafting the manuscript. MA-MM cleaned the sequence data. RN provided the laboratory facility, reagents and sequence analysis. All authors read and approved the final manuscript.

## Acknowledgments

We are grateful to the animal owners and Atbara Butana Station, UmBanein Kenana Station, Kuku Research Center and Multi-Purposes Company for allowing us to sample their animals.

